# Pan-cancer analysis of transcriptional metabolic dysregulation using The Cancer Genome Atlas

**DOI:** 10.1101/295733

**Authors:** SR Rosario, MD Long, HC Affronti, AM Rowsam, KH Eng, DJ Smiraglia

## Abstract

Understanding the levels of metabolic dysregulation in different disease settings is vital for the safe and effective incorporation of metabolism-targeted therapeutics in the clinic. Using transcriptomic data from 10,704 tumor and normal samples from The Cancer Genome Atlas, across 26 disease sites, we developed a novel bioinformatics pipeline that distinguishes tumor from normal tissues, based on differential gene expression for 114 metabolic pathways. This pathway dysregulation was confirmed in separate patient populations, further demonstrating the robustness of this approach. A bootstrapping simulation was then applied to assess whether these alterations were biologically meaningful, rather than expected by chance. We provide distinct examples of the types of analysis that can be accomplished with this tool to understand cancer specific metabolic dysregulation, highlighting novel pathways of interest in both common and rare disease sites. Utilizing a pathway mapping approach to understand patterns of metabolic flux, differential drug sensitivity, can accurately be predicted. Further, the identification of Master Metabolic Transcriptional Regulators, whose expression was highly correlated with pathway gene expression, explains why metabolic differences exist in different disease sites. We demonstrate these also have the ability to segregate patient populations and predict responders to different metabolism-targeted therapeutics.

## Introduction

Despite waning interest in how metabolism influences cancer, recent efforts have brought a renewed awareness of cancer as a metabolic disorder^1-4^. While the field was first introduced to cancer as a glycolytic disease, often described as the Warburg Effect^5^, modern advancements and technologies have pointed to other metabolic dependencies, such as fatty acid metabolism in Prostate Cancer^6^. These recent investigations have led to the inclusion of metabolic reprogramming as a new hallmark of malignant transformation^7^. However, the extent to which all metabolic genes and pathways are expressed by cancers of different origins, and how they differ from one another, is largely underexplored. Even more pressingly, how these metabolic pathways and genes differ from the non-malignant, normal human tissues has yet to be determined. Few papers exist that attempt to explain differences in cancer and normal that can be leveraged to understand metabolic reprogramming, based on genomic perturbation^8-11^. While some innate differences between tumor tissues and normal are addressed, the focus is typically on common mechanisms and metabolic gene dysregulation that exist pan-cancer, rather than those changes that exist in an individual tumor type at the transcriptomic level, and how these affect existing chemotherapeutic treatment^8,10,11^. Others look at this concept solely from the metabolomics angle, in a single disease site^9^. These studies, while informative, create a gap in which we question if there are targetable metabolic pathways unique to a single disease site, or globally dysregulated. Additionally, we can question whether there is a way to distinguish those patients that will respond to these metabolic-targeted therapies, based on their distorted metabolism.

Scientific consortiums like The Cancer Genome Atlas (TCGA)^12^ encourage comprehensive genomics approaches in large numbers of patients with many different cancer types, as well as their matched normal tissues. These data allow the opportunity to address whether metabolic genes differ between normal and malignant conditions across diverse tissues of origin. While metabolomics, the systemic study of the small molecules utilized and left behind during essential cellular processes^13^, is the most comprehensive way to understand the metabolic composition of a cell at a given time, the technique is still in its infancy^14^. Conversely, abundant and readily available transcriptomic data exist for a large number of patients in many types of cancer. Such data sets provide the opportunity to investigate the variety of mechanisms cancers utilize to control metabolic enzyme expression in order to achieve metabolic reprogramming, including feedback and crosstalk between metabolite pools and transcription^15-16^.

Recently, transcriptomics data in conjunction with current biochemical understanding have been exploited to construct genome-scale metabolic workflows^17^. For instance, the metabolic output in *E.coli* was successfully predicted using transcriptomic data, in which over half of the metabolic outputs from greater than 450 different reactions within the organism were correctly modeled^18-21^. Many different algorithms exist that attempt to extrapolate metabolic output from transcriptomic inputs in model organisms, with varying accuracy and sensitivity^22-23^. Nevertheless, extrapolating metabolic changes from transcriptomics is not without its challenges, as stoichiometric relationships and kinetic information must be assumed in many cases^18^. Recently, evidence for a high level of significant correlation between gene expression and metabolite levels was revealed in a detailed look at breast cancer RNA-sequencing and unbiased metabolomics^13^.

An additional challenge to understanding metabolic reprogramming in cancer, lies in determining the genetic and epigenetic changes that control the metabolic phenotypes. To this end, we suggest that elucidating expression and alteration of Master Metabolic Transcriptional Regulators (MMTRs) may provide novel understanding of why metabolism differs in varying tissues and provide new targets for the treatment of tumors. Susumu Ohno first recognized master transcriptional regulators in the developmental field in the 1970s^24^, using the term to describe transcription factors that regulate sets of genes that determine developmental fate. Master regulators (MRs) have been implicated in a variety of disease states^25-27^ and with several genomic alterations^28-29^. More recently, MRs have become interesting as biomarkers of disease^30-31^ and also as pharmacological targets^32^, with the intention that targeting the MR would also modulate the downstream targets.

Master regulators have also been shown to exert regulatory control over specific metabolic pathways. One example is the sterol regulatory element binding proteins (SREBPs) that control lipid metabolism. This family of genes is highly associated with the expression of genes and enzymes involved in cholesterol, fatty acid, triacylglycerol and phospholipid synthesis^33^. Transgenic and knockout mice have further elucidated the effect of these master regulators on lipid metabolism, showing that the unique regulation and activation properties of each of the three SREBP isoforms facilitate homeostatic regulation of lipid metabolism^34-35^. However, a more nuanced understanding of unique metabolic dependencies, or weaknesses, in specific cancer types, subtypes, or even tissues of origin, may provide novel therapeutic targets that have better potential for lower toxicity than traditional chemotherapeutics. A recent example is the recognition that cancers that are deficient in the methylthioadenosine phosphorylase (MTAP) enzyme are highly susceptible to inhibition of methionine adenosyl transferase 2A (MAT2A) resulting in reduced function of protein methyltransferase 5 (PRMT5)^36^. Metabolic therapies provide an attractive approach in the clinic, as it has not only evaded drug resistance thus far, but has also been shown to prevent multi-drug resistance in tumors^37^. Determining responders to these metabolic therapies, however, provides a challenge. There are currently no studies determining the master transcriptional metabolic regulators (MMTR) of specific metabolic pathways, which may serve as drivers of metabolic phenotypes. These would also provide new insights into ways to therapeutically leverage metabolic dependencies and segregate patient populations in terms of response to metabolic-targeted therapeutics.

The aim of this study was to comprehensively assess which metabolic pathways have altered transcriptional profiles in 26 different cancer types as compared to their matched normal tissues. This information was then utilized to identify those pathways that were uniquely altered in malignancies from specific tissues of origin or those that were commonly altered across all cancer types, as well as those metabolic pathways that were most altered in specific molecular subtypes that exist within a cancer type. Here we demonstrate that we have the ability to not only segregate different disease sites and different molecular subtypes of the same disease, but also to predict response to metabolism-targeted therapy. This selective drug sensitivity is further explained by MMTRs we identified for the individual pathways. This represents a means of identifying a mechanism by which these metabolic pathways become distorted in malignancy and to offer novel targets for intervention.

## Results

### Pan-cancer screen for transcriptional metabolic dysregulation

To screen for transcriptional metabolic dysregulation, we used RNA-sequencing data from 26 different types of cancer with matched normal samples from TCGA^12^ (Fig 1A, Supplementary Table 1) using a custom analysis pipeline as depicted in Figure 1. Magnitude of metabolic dysregulation was calculated by first determining a list of Differentially Expressed Genes (DEGs), which includes log fold changes and adjusted p-values, comparing tumor tissues with normal matched samples and assigning scores based on 114 metabolic pathways sourced from The Kyoto Encyclopedia of Genes and Genomes (KEGG) ^38^. Adjusted p-value magnitude is affected by sample size, which varies across data sets. To account for how such variation would affect the metabolic pathway score, each was divided by the square root of n (the sample size) for each disease site. Bootstrapping methods were then used to determine which pathways were significantly dysregulated (red) or non-significantly dysregulated (gray) (Fig 1B). The metabolic dysregulation scores (Fig 1C) were then confirmed in separate patient populations for prostate^39^, lung adenocarcinoma^40^, and breast carcinoma^41^ (Supplementary Figure 1). Pathway scores in validation cohorts were significantly correlated to those generated in TCGA cohorts from the same disease sites, demonstrating the robustness of the pipeline. To determine if there were patterns within the classes of metabolic dysregulation, individual pathway scores were segregated into major classes of metabolism (Fig 1D). MMTRs were then determined for individual pathways, such as the Pentose Glucuronate Interconversion Pathway (Fig 1E), as a means of elucidating the drivers of unique metabolic phenotypes that exist in cancers of different origins.

**Figure 1.**
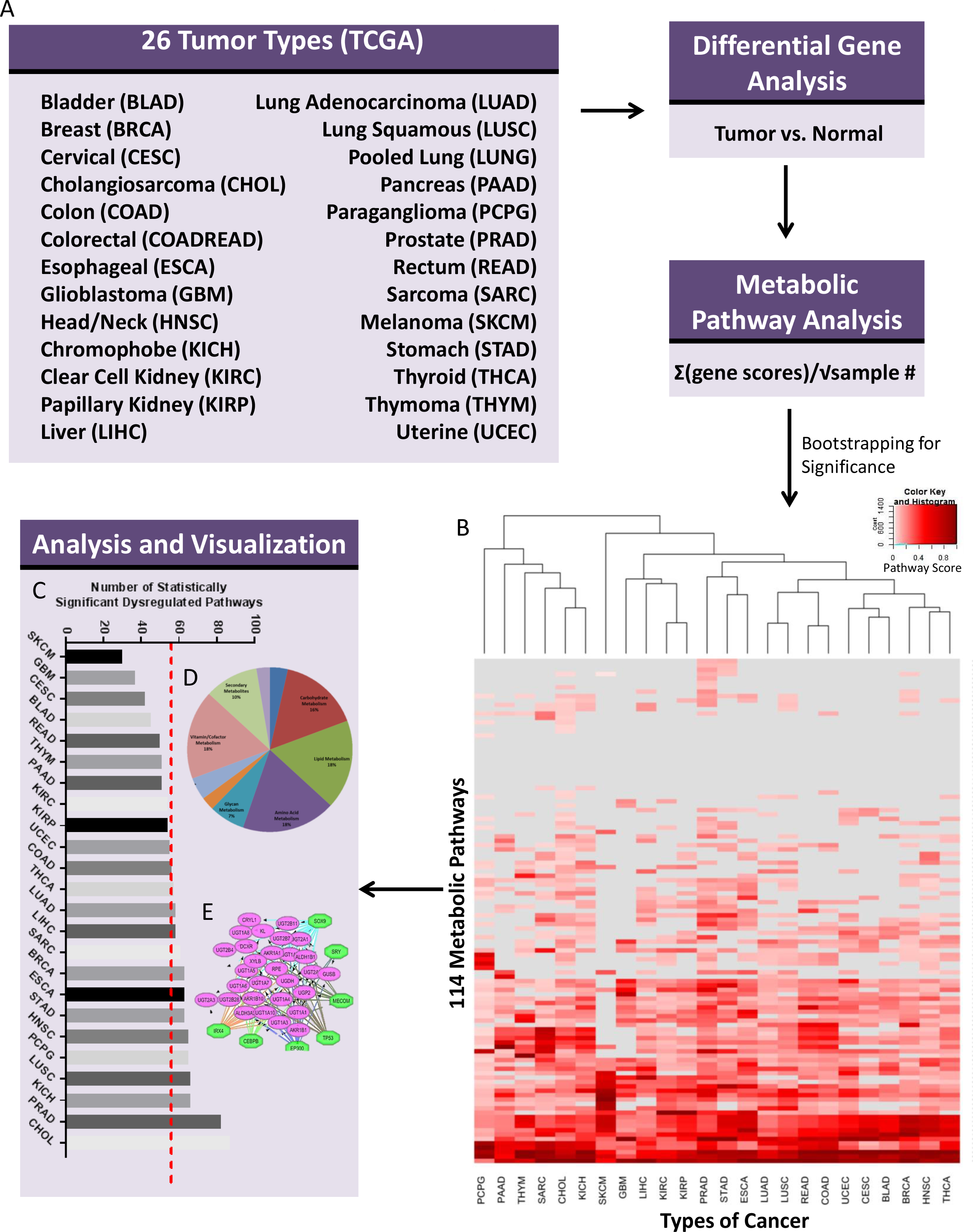
Transcriptional Metabolic Pathway Analysis Methods Pipeline. (A) 26 cohorts of tumor samples, including two pooled sets (COADREAD and LUNG) from The Cancer Genome Atlas (TCGA), with matched Normal Samples, were utilized to determine the transcriptional metabolic profiles specific to each type of cancer, as compared to their normal. (B) Pathway scores ((ΣlogFC*-log(adj.p.val))/√n), for 114 Metabolic Pathways from KEGG, were then calculated based on the results of Differential Expressed Gene (DEG) analysis using Limma to compare tumors to matched normal. Pathways are then bootstrapped for significance, to determine which pathways are highly dysregulated as compared to chance. Those pathways are then plotted in a heatmap, with the type of cancer as the X-axis and the 114 pathways as the y-axis. Pathways that are non-significant are gray and a gradient from white to red for those pathways significantly dysregulated and the intensity of red indicating the magnitude of dysregulation. (C) Significant pathway scores are then summed to determine with of the types of cancers are most metabolically dysregulated at the transcriptional level. (D) The pathways were then sorted into each of the 10 major metabolic pathway subtypes defined by KEGG and later underwent (E) Master Regulator Analysis via iRegulon.

The 114 individual metabolic pathways were subsequently condensed into ten major categories of metabolism based on KEGG classifications (Fig 2A). After bootstrapping, pathways in each classification were then further broken down by the number of cancers for which they were dysregulated (Fig 2B). Additionally, we identified unique pathways, which were altered in just one disease site (Fig 2C). For example, Prostate cancer (PRAD) had two pathways uniquely dysregulated (Polyamine Biosynthesis and Nicotinamide Adenine Dinucleotide Biosynthesis). Further, analysis of pathways within the major metabolic categories revealed patterns of dysregulation reflective of known common metabolic reprogramming in cancer. For example, within the carbohydrate metabolism category we found patterns consistent with the Warburg Effect, such as dysregulation of glycolysis and gluconeogenesis (Fig 2D), to varying degrees in all cancer types. While these results support those found in the literature, the method also allowed for novel observations regarding metabolic disruption across the 26 types of cancer examined.

**Figure 2.**
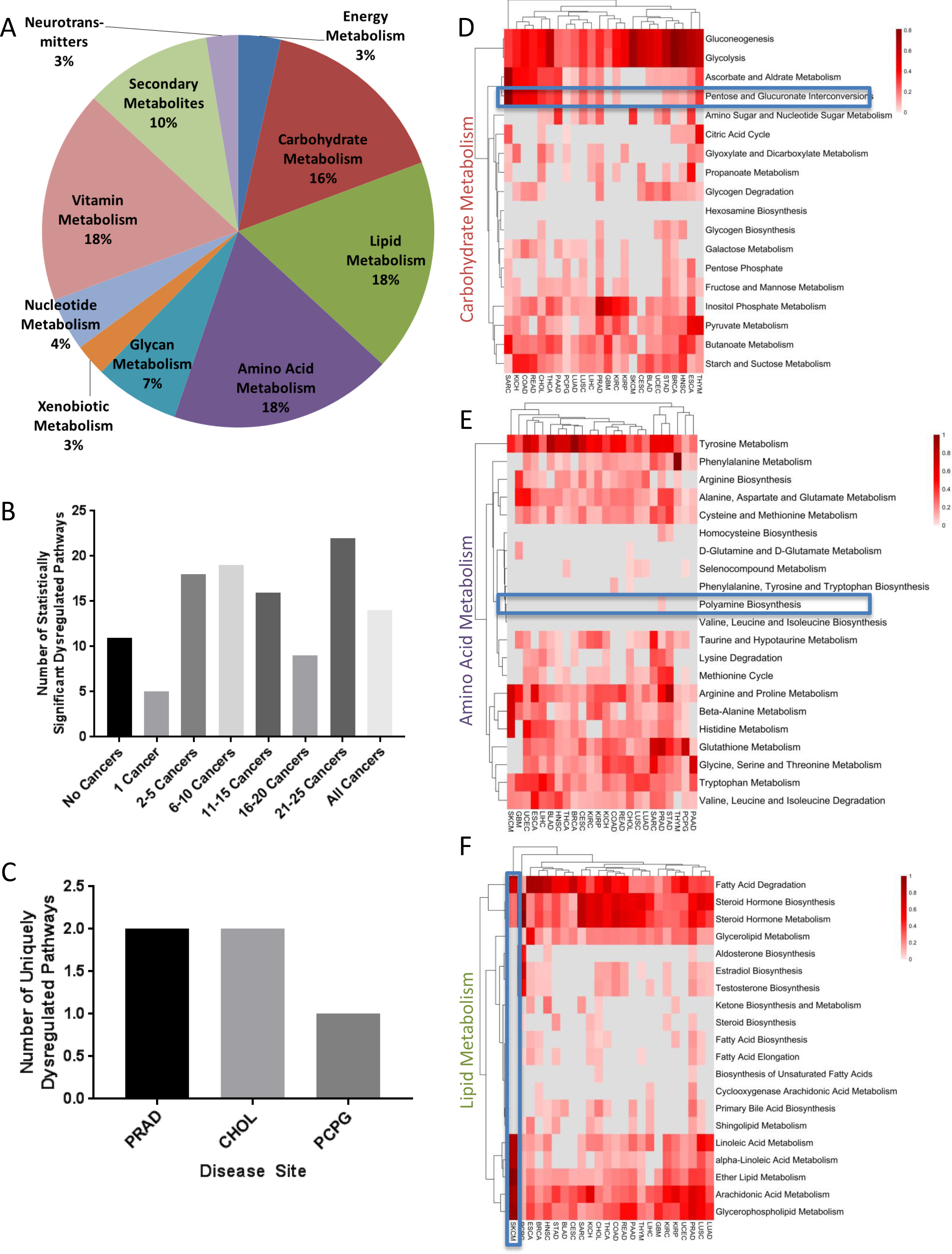
Ascertaining Pathways of interest by looking at major types of metabolic pathways. (A) 114 Metabolic KEGG Pathways broke down into 10 major metabolic types of pathways. This allowed for the identification of (B) pathways that were statistically significantly altered in a variety of cancers and (C) the number of uniquely dysregulated pathways in specific tumor types. Each of these major metabolic categories was then broken down into individual heatmaps of bootstrapped pathway scores where gray are non-significant altered pathways and the gradient of red represents the magnitude of dysregulation in each of the pathways across the cancer cohorts. (D) In Carbohydrate Metabolism, a pathway largely altered across all tumor types, like Pentose and Glucuronate Interconversions, is highlighted. (E) Within Amino Acid Metabolism the Polyamine Biosynthetic Pathway is highlighted as an example of cancer type specific dysregulation. (F) Subcutaneous Melanoma (SKCM) was identified as the cancer type with the highest degree of dysregulation, based on Euclidian distance, within a subset of the KEGG pathways in the Lipid Metabolism category.

One such finding is the common, but not universal, dysregulation of the Pentose and Glucuronate Interconversion Pathway (Fig 2D), which is significantly altered in 20 cancer types. Due to its high degree of dysregulation, this pathway has been studied in some cancer types, including LIHC^42-43^. However, little has been done to study this pathway in SARC, where we find it is most strongly dysregulated. Meanwhile in only six types of cancer (Uterine Corpus Endometrial Carcinoma (UCEC), Bladder Urothelial Carcinoma (BLAD), Cervical and Endocervical cancers (CESC), Skin Cutaneous Melanoma (SKMC), Kidney Renal Papillary Cell Carcinoma (KIRP), and Glioblastoma Multiforme (GBM)) the Pentose Glucuronate Interconversion Pathway was not significantly dysregulated. Conversely, some pathways were only dysregulated in a single type of cancer. For instance, the Polyamine Biosynthetic Pathway was only significantly dysregulated in prostate cancer (PRAD), suggesting unique dependencies of certain cancers (Figure 2E).

In addition, within specific metabolic categories, hierarchical clustering highlighted disease sites with distinct levels of dysregulation. An example of this is the disruption of Lipid metabolism in Skin Cutaneous Melanoma (SKCM), whose level and pattern of dysregulation among pathways within this category caused this disease site to segregate separately from others (Fig 2F). SKCM is one of the least metabolically dysregulated cancer types, in terms of the number of significantly dysregulated pathways (Fig 1B). As shown in Figure 1C, among the 26 cancer types, the median number of significantly dysregulated pathways is 58 out of the 114, while for SKCM there are only 30.

### Pathways dysregulated across multiple cancer subtypes

The Pentose Glucuronate Interconversion pathway clusters closely with the Glycolysis and Gluconeogenesis pathways that are significantly dysregulated in all cancer types (Fig 2B). This pathway is involved in the interconversions of monosaccharide pentose and glucuronates, which are the salts or esters of glucuronic acid^41^. While it is known that this pathway is frequently dysregulated in hepatocellular carcinoma, little is known about the dysregulation of this pathway in the context of other disease sites^42-43^. As is shown in the heatmap of carbohydrate metabolic pathways, many cancer types highly dysregulate this pathway, while only a few do not (Fig 2B). To better understand the expression changes within the Pentose Glucuronate Interconversion pathway, the 35 individual genes that make up the pathway were examined in detail across cancer types (Figure 3A-B). This approach demonstrated that there are two distinct groups of cancer sites, those that significantly up-regulate many of the genes within this pathway, and conversely those that down-regulate a majority of genes within this pathway.

**Figure 3.**
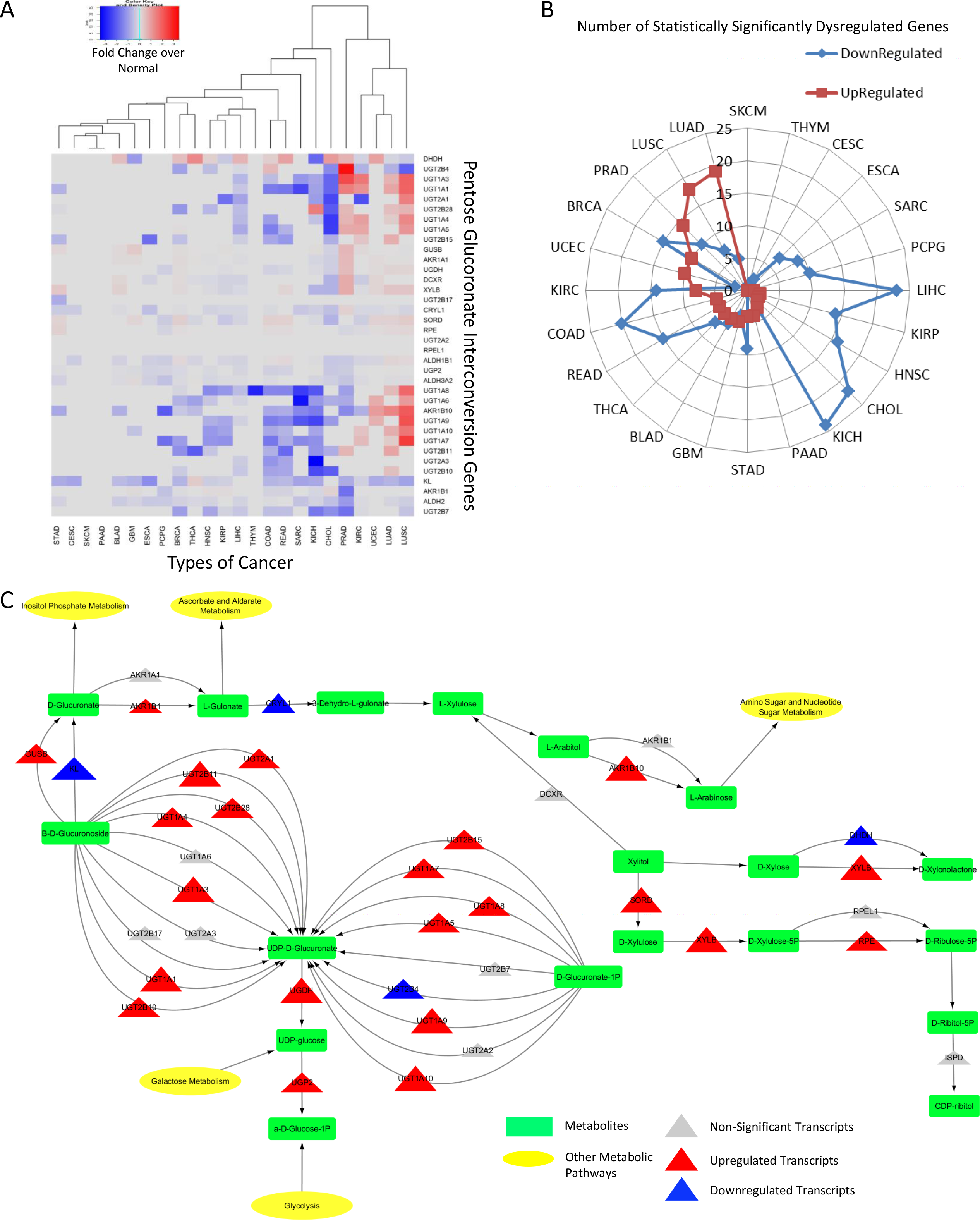
The Pentose Glucuronate Interconversion Pathway is dysregulated across all types of cancer but to a different extent and in different directions. (A) Heatmap of fold changes, showing the direction of transcriptional change in tumor over normal for each individual gene within the Pentose Glucuronate Interconversion Pathway, across all cancer types. (B) Radar plot demonstrating the number of statistically significant up-regulated (red) and down-regulated (blue) genes within a pathway, for each of the indicated cancers. (C) Model of the Pentose Glucuronate Interconversion Pathway in LUAD, which has the largest number of statistically significant up-regulated genes. Pathway displays genes (triangles) shaded by direction (red and blue, up and down, respectively) and significance and sized by fold change differences. Pathways also include metabolite outputs (green rectangles) and connected pathways (yellow ellipse).

Consistent with what has previously been reported in LIHC, our analysis found this disease site to be among the cancers with the most significantly dysregulated genes, nearly all of which are down-regulated (23 out of 35). In addition we made the novel observations that SARC, Kidney Chromophobe (KICH), Cholangiocarcinoma (CHOL), and Colon Adenocarcinoma (COAD), also have a high degree of down-regulation within the Pentose Glucuronate Interconversion pathway but the magnitude of down-regulation exceeds that of LIHC. Conversely, the lung cancer subtypes, lung adenocarcinoma (LUAD) and lung squamous cell carcinoma (LUSC), up-regulate the greatest number of genes within the pathway, though the magnitude of upregulation is far greater in LUSC. While it has been reported that intermediate metabolites of the Pentose Glucuronate Interconversion pathway are increased in LUAD^44^, the transcriptional upregulation of 20 of the 35 genes within this pathway are unknown. Previous literature has also pointed to the dysregulation of this pathway in LUSC but failed to further explore how this pathway was transcriptionally disrupted^45^.

Modeling the metabolic pathways by placing the significantly dysregulated genes within the context of the metabolic circuit, may predict which metabolites will be most readily affected and in what direction. For example, comparing the Pentose Glucuronate Interconversion pathway models in LUAD (Fig 3C) and LIHC (Sup Fig 2), reveal two different metabolic pictures. A large number of the genes within the pathway, that are upregulated in LUAD, but downregulated in LIHC, contribute to the generation of UDP-D-Glucuronate from β-D-Glucuronoside and D-Glucuronate-1 phosphate. Therefore, the expression levels of these enzymes predicts for relatively high levels of UDP-D-Glucuronate in LUAD, but low levels in LIHC. This finding is highly consistent with previous metabolomics studies in LUAD, which have asserted that there is an increased level of UDP-D-Glucuronate, in particular, in cancer tissues, as compared to matched normal^44^, as well as the literature regarding a global down regulation of metabolites within the Pentose Glucuronate Interconversion Pathway in LIHC^46^.

### Pathways uniquely dysregulated within a single cancer subtype

This metabolic pipeline can also elucidate pathways that are uniquely dysregulated in a specific cancer type. An example of this is the unique disruption of the Polyamine Biosynthetic Pathway in PRAD (Fig 2E, Fig 4A). Polyamines are small, positively charged molecules with a multitude of functions, impacting almost every aspect of cell survival^47^. While this pathway is important in every cancer, the unique dysregulation of this pathway in PRAD is of particular interest because flux through the biosynthetic pathway is already extremely high in normal prostate due to the high rate of secretion of acetylated polyamines to the prostatic lumen^48^. Not only does PRAD have the highest number of significantly upregulated (as compared to normal prostate) genes among the 13 assigned to this pathway (Fig 4B), they also show the greatest magnitude of change, which is reflected in the large Euclidean distance observed in the unsupervised hierarchical clustering (Fig 4A).

**Figure 4.**
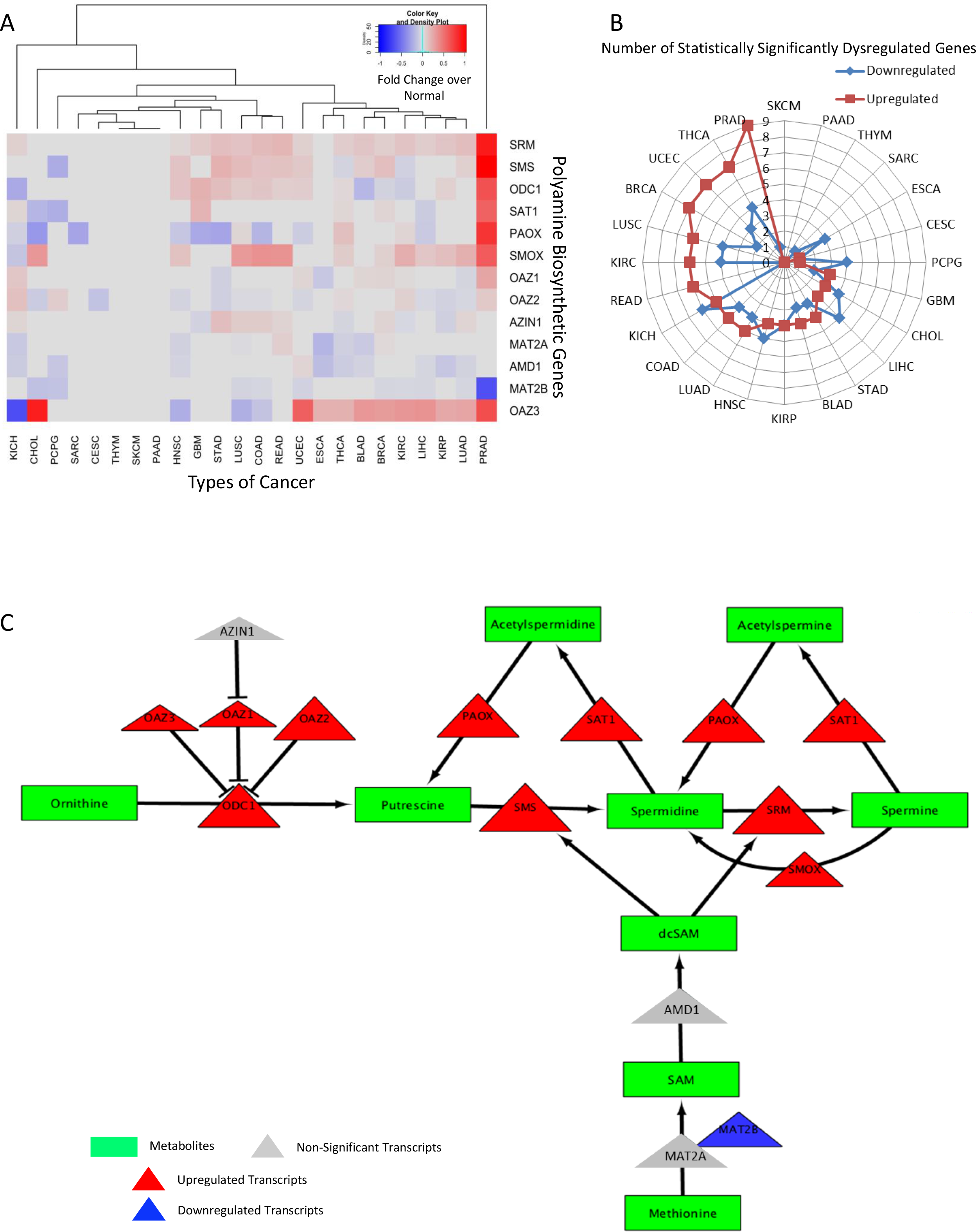
The Polyamine Biosynthetic Pathway is highly specifically dysregulated in Prostate Cancer. (A) Heatmap of fold changes, showing the direction of transcriptional change in tumor over normal for each individual gene within the Polyamine Biosynthetic Pathway, across all cancer types. (B) Radar plot demonstrating the number of statistically significant up-regulated (red) and down-regulated (blue) genes within a pathway, for each of the indicated cancers. (C) Model of the Polyamine Biosynthetic Pathway in PRAD, which has the largest number of statistically significant up-regulated genes. Pathway displays genes (triangles) shaded by direction (red and blue, up and down, respectively) and significance and sized by fold change differences. Pathways also include metabolite outputs (green rectangles).

Modeling of the polyamine metabolic circuit clearly demonstrates an increase in polyamine biosynthesis and catabolism in PRAD (Fig 4C). This is reflected in increased expression of both rate limiting enzymes in the biosynthetic pathway (*ODC1* and *AMD1*), as well as significant increases in the catabolic enzymes *SAT1, PAOX* and *SMOX*. The pathway is further enhanced by upregulation of *MAT2A* and down regulation of the inhibitory subunit *MAT2B*, predicting for enhanced SAM production feeding into *AMD1* as well as greatly increased expression of the polyamine synthases, *SRM* and *SMS*. These findings are consistent with the well-documented increased level of polyamines, acetylated polyamines, and other metabolites within the Polyamine Biosynthetic pathway in prostate cancer, all of which can be predicted by the modeling approach^49-50^. Conversely, in KICH, the two rate limiting enzymes *ODC1* and *AMD1* are significantly down regulated, suggesting reduced levels of polyamines (Fig 4A, Sup Fig 3). Additionally, there is an increase in *SAT1*, which leads to acetylation of the polyamines, but a decrease in *PAOX*, which leads to decreased back conversion of acetylated polyamines to un-acetylated polyamines. All of these transcriptional changes, taken together, lead to the depletion of polyamines in the cancer state, as compared to the normal tissues. This broad upregulation of the Polyamine Biosynthetic Pathway in PRAD would suggest a unique dependence on its function, as compared to other disease sites, providing rationale for pharmacological intervention.

The drug N^1^, N^11-^bis(ethyl)norspermine (BENSpm), an SAT1 stabilizer that increases polyamine acetylation, was utilized to understand whether prostate cancer cell lines could more selectively be targeted by further destabilization of polyamine biosynthesis. When comparing the sensitivity of seven cell lines, two prostate cancer (DU145 and PC-3), two kidney cancer (ACHN and 786-O), and three breast cancer cell lines (MDA-MB-231, HS578T and MCF7) (Fig 5A, 5B), the prostate cancer lines were the most sensitive to BENSpm treatment compared to any other cell line.

**Figure 5.**
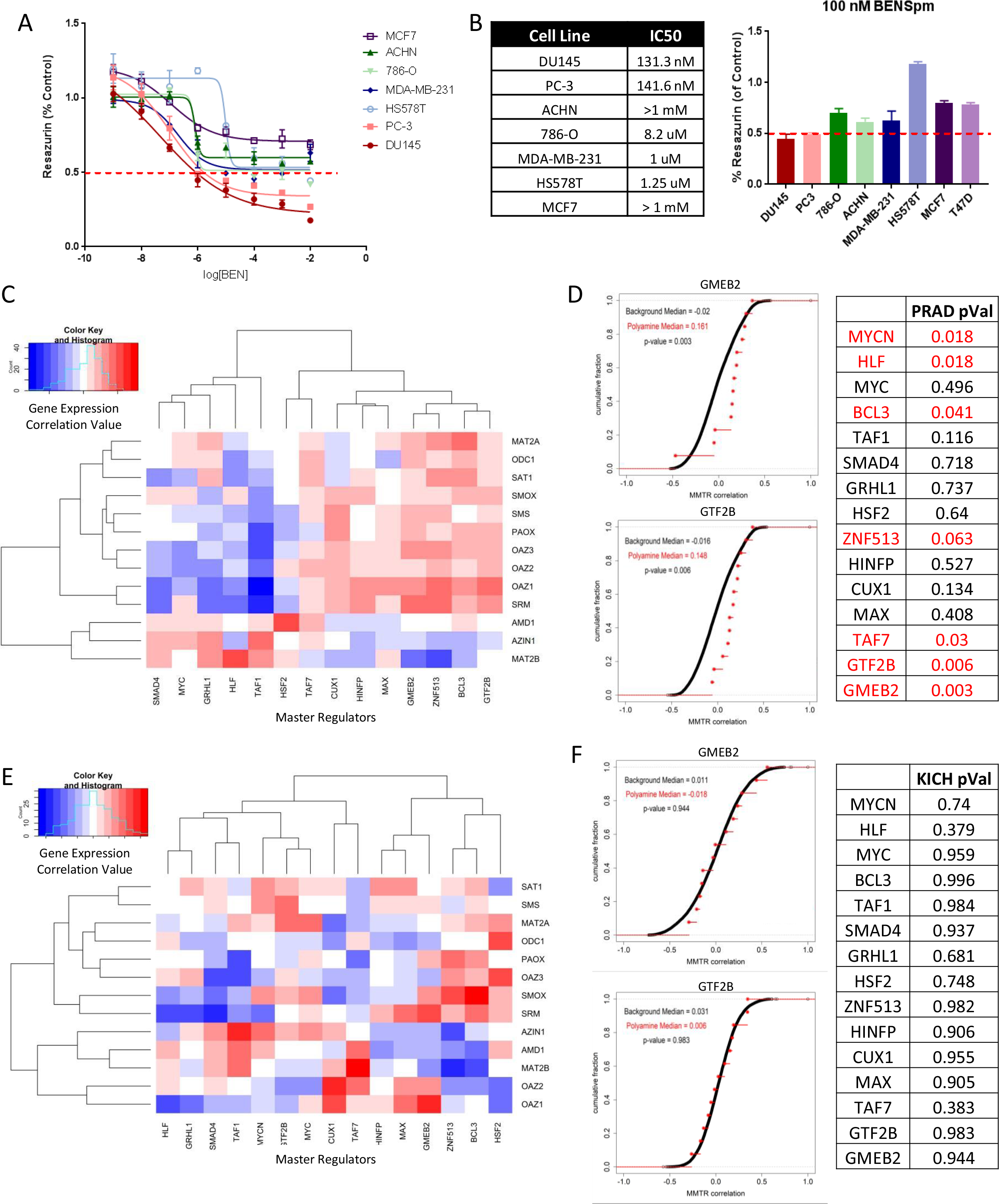
Targeting Polyamine Biosynthesis is highly effective in Prostate Cancer and Master Metabolic Transcription Regulators (MMTRs) may explain why. (A) Dose response curves show prostate cancer cell lines (PC3 and DU145) are most sensitive to SAT1 activation by BENSpm, as compared to kidney cancer cell lines (ACHN and 786-O) and breast cancer cell lines (MCF7, MDA-MB-231, and HS578T). (B) This is confirmed by calculation of IC50s. (C) Prostate Cancer (PRAD) heatmap of correlations between Master Metabolic Transcription Regulators (x-axis) and the Polyamine Biosynthetic Genes (y-axis) with the intensity representing strength of correlation (blue is negative, red is positive)and (D) Cumulative Distribution frequencies showing MMTR correlation with every gene in the genome, with red dots indicating the correlations with Polyamine Biosynthetic genes, show a distinct pattern and high level of statistically significant correlation values between MMTRs and Polyamine Biosynthetic genes. P-values for all MMTRs and Polyamine genes are reported in the table, with significant associations highlighted in red (E) Kidney Chromophobe (KICH) heatmap of correlations between Master Metabolic Transcription Regulators (x-axis) and the Polyamine Biosynthetic Genes (y-axis) with the intensity representing strength of correlation (blue is negative, red is positive)and (F) Cumulative Distribution frequencies showing MMTR correlation with every gene in the genome, with red dots indicating the correlations with Polyamine Biosynthetic genes, show an random pattern and non-statistically significant correlation values between MMTRs and Polyamine Biosynthetic genes. P-values for all MMTRs and Polyamine genes are reported in the table, with significant associations highlighted in red

Master Metabolic Transcriptional Regulators (MMTRs) may explain these differences in polyamine biosynthesis metabolism dysregulation between cancer types. When analyzing the overall pathway scores, the two most highly dysregulated cancer types are PRAD and KICH (Fig 4A). As previously demonstrated, when directionality is taken into account, these pathways are largely dysregulated in opposite directions (Fig 4A). Utilizing iRegulon^51^, which pairs motifs and ChIP-Seq tracks to determine transcription factors that control the expression of gene networks, a list of master transcriptional regulators of the polyamine biosynthetic pathway was constructed (Sup Fig 4). We then correlated expression of MMTRs with the expression of each of the individual genes in the Polyamine Biosynthetic Pathway in both PRAD and KICH cohorts (Fig 5C). Distinct patterns of correlation emerged in PRAD, where a majority of the MMTRs were significantly correlated, either positively or negatively, with Polyamine Biosynthetic genes. Comparisons of the cumulative distribution frequencies of correlations between MMTRs and all genes in the genome (black) with correlations of these MMTRs with polyamine biosynthetic genes only (red) show that polyamine gene correlations are statistically significant when considering global expression patterns, demonstrated by a shift in the distributions (Fig 5D). Conversely, KICH lacks a strong pattern of correlation between MMTRs and Polyamine Biosynthetic genes (Fig 5E). This finding was confirmed by cumulative distribution analysis, where there was no significant relationship between MMTR and Polyamine Biosynthetic gene expression observed when considering global transcriptional patterns (Fig 5F). The top four MMTRs(BCL3, GMEB2, GTF2B, and ZNF513), with the strongest collective positive correlation, are important for regulation of polyamine biosynthetic genes, as evidenced by an iRegulon network, which demonstrates these 4 MMTRs are collectively predicted to regulate 10/13 of the genes from the pathway (Sup Fig 4). Thus, the metabolic analysis pipeline can accurately provide possible novel targets of pharmacologic intervention. Further, this information combined with master regulator analysis, can be a useful tool providing new insights into drivers of metabolic reprogramming across and within cancer sites.

### BRCA Subtype Metabolic Reprogramming

BRCA is one of the most metabolically dysregulated cancer types, in terms of the sum of pathway scores (Fig 1C). Analysis of all BRCA cases reveals a large level of dysregulation of Carbohydrate, Lipid, and Amino Acid Metabolism, in roughly equal proportions (Fig 6A). Additionally, the top dysregulated pathways seem to be a mix of these larger categories, encompassing pathways from each major category (Fig 6B). Importantly, BRCA consists of 4 major molecular subtypes with distinct treatments and outcomes for patients: Luminal A, Luminal B, HER2 and Basal^52^. This, therefore, lead us to question whether these 4 major subtypes had distinct transcriptional metabolic profiles.

**Figure 6.**
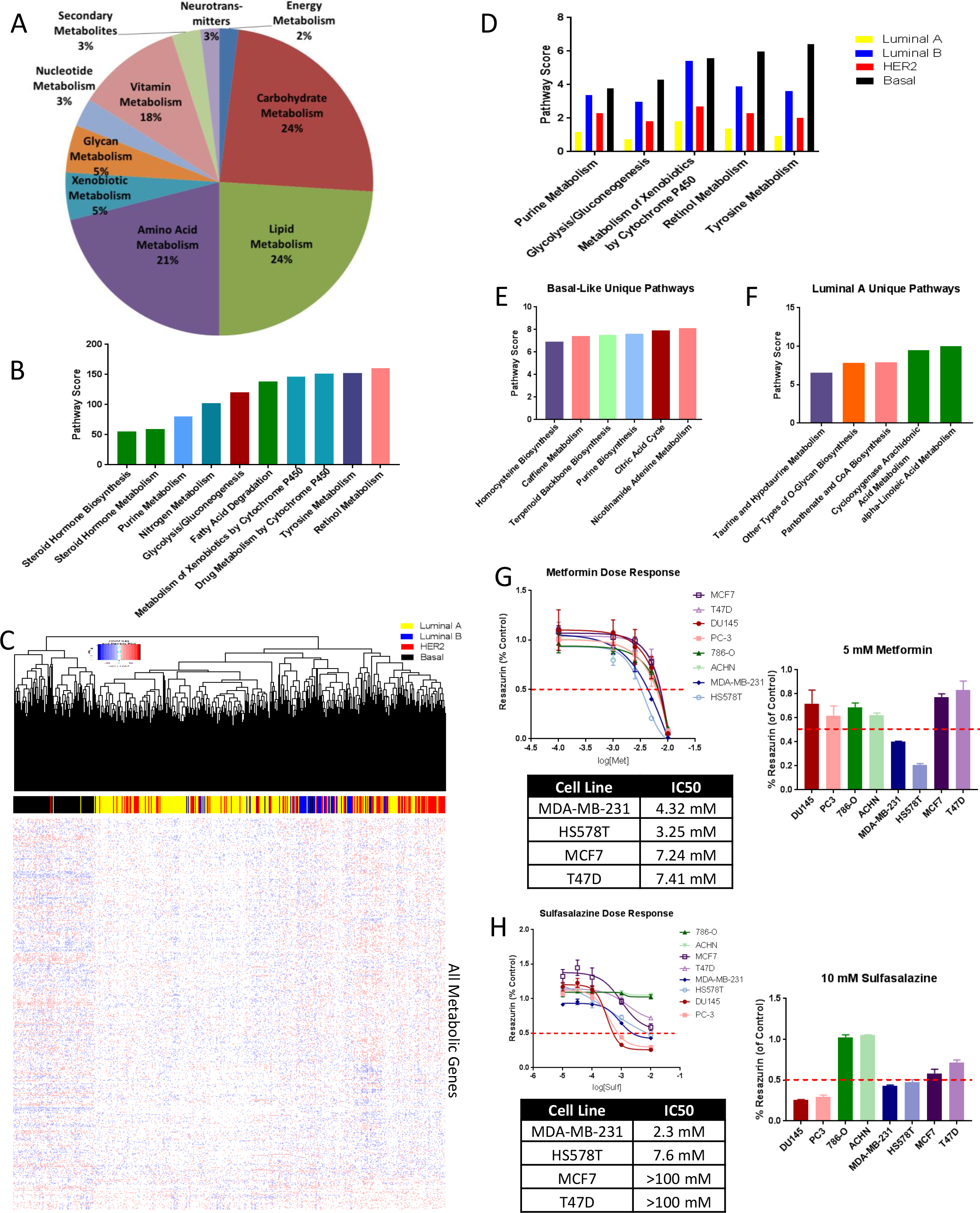
Metabolic dysregulation distinguishes BRCA molecular subtypes and defines therapeutic sensitivity. (A) Breakdown of 62 significantly dysregulated metabolic pathways in the BRCA pooled data. There is a roughly equal dysregulation of amino acid, carbohydrate and lipid associated pathways. (B) The top pathways dysregulated in BRCA come from a wide variety of major categories, like nucleotides and lipids. However, the two most dysregulated pathways in BRCA are a vitamin-associated pathway, Retinol Metabolism, and an amino acid pathway, Tyrosine Metabolism. (C) Unsupervised clustering of BRCA patients, who were classified based upon the PAM50, on all metabolic genes, reveals a tight cluster of the Basal-like subtype (black), which are highly metabolically dysregulated as explained by the magnitude of dysregulation, as displayed in the heatmap. While Luminal A (yellow), Luminal B (blue) and HER2-expressing (red), did not cluster as tightly, there are still recognizable groups of these patients. (D) The Top 5 pathways that overlapped between all 4 subtypes of patients are shown here. While these pathways are highly dysregulated in all 4 subtypes, they vary to different extents and are almost always highest in the Basal-like cells. (E) After metabolic pathway scoring, pathways unique to the Basal-like, most aggressive, and (F) Luminal A most indolent, patients were isolated and plotted. The top 5 unique pathways for each of the subtypes are shown. (G) Targeting the Citric Acid Cycle with Metformin revealed increased sensitivity in Basal-like cells, as compared to Luminal A cells, as emphasized by the IC50 values. (H) Targeting the Homocysteine Biosynthetic Pathway with Sulfasalazine revealed increased sensitivity in Basal-like cells, as compared to Luminal A cells, as emphasized by the IC50 values.

Using the PAM50^53, 54^, a set of 50 genes whose differential expression is utilized to classify BRCA, all patients were assigned to one of the four subtypes. Patients were first randomly clustered based on the expression of all metabolic genes (Fig 6C). Basal-like tumors (black), clustered out very distinctly from the Luminal A (yellow), Luminal B (blue), and HER2 expressing (red) counterparts, indicating a strong shift of metabolic phenotype in these patients. While not as distinct, smaller clusters did form for each of the other molecular subtypes. Each molecular subtype was then compared to the normal tissues, and DEG lists for each independent cluster was utilized to determine which of the 114 pathways were significantly dysregulated. This analysis revealed a total of 89 dysregulated pathways, some of which were missed entirely by an analysis of the BRCA pooled data. Pathways like Tyrosine Metabolism and Retinol Metabolism, as well as Glycolysis and Gluconeogenesis are dysregulated across all subtypes, but to a different degree (Fig 6D), as indicated by the different scores for each individual pathway among the four molecular subtypes. While many pathways were dysregulated across all subtypes, there were distinct pathways present in each subtype of the disease. For example, in Basal-like tumors, the most aggressive form of BRCA, the Terpenoid Backbone Biosynthetic Pathway, Homocysteine Biosynthesis and the Citric Acid Cycle are uniquely dysregulated, amongst others (Fig 6E). Meanwhile, in Luminal A tumors, the subtype with the most favorable prognosis, we found that alpha-Linoleic Acid Metabolism, Taurine and Hypotaurine Metabolism and Cyclooxygenase Arachidonic Acid Metabolism were uniquely dysregulated (Fig 6F). Further, this dysregulation of unique metabolic pathways in molecular breast subtypes would suggest a potential differential sensitivity to therapeutic intervention.

The drug Metformin, a first generation biguanide that decreases glucose metabolism through the Citric Acid Cycle^55^, was utilized to understand whether basal breast cancer cell lines could more selectively be targeted. When comparing the sensitivity of eight cell lines, two prostate cancer (DU145 and PC-3), two kidney cancer (ACHN and 786-O), two luminal breast cancer cell lines (MCF7 and T47D) and two basal breast cancer cell lines (MDA-MB-231 and HS578T) (Fig 6G), we did in fact see increased sensitivity of basal breast cancer cell lines to Metformin, as compared to any other cell line. Additionally, because Homocysteine Biosynthesis is specifically dysregulated in the basal like subtype, we utilized the drug Sulfasalazine, an Xc cysteine-glutamate transport inhibitor that decreases intracellular homocysteine pools^56^, to understand whether basal breast cancer cell lines could more selectively be targeted by destabilization of the Homocysteine Cycle. When comparing the sensitivity of eight cell lines, two prostate cancer (DU145 and PC-3), two kidney cancer (ACHN and 786-O), two luminal breast cancer cell lines (MCF7 and T47D) and two basal breast cancer cell lines (MDA-MB-231 and HS578T) (Fig 6H), we did in fact see increased sensitivity of basal breast cancer cell lines to Sulfasalazine, as compared to the luminal breast cancer cell lines. Interestingly, prostate cancer cell lines were even more sensitive, which is a finding consistent with the fact that our analysis identifies PRAD as one of two disease sites with significantly dysregulated Homocysteine Biosynthesis (Fig 2E). It is also notable that this pathway was not detected as dysregulated in the BRCA cohort, but it is dysregulated specifically in the basal like subtype.

Using the genes from both the unique and overlapping pathways, MMTRs of both the uniquely dysregulated pathways in each subtype (Fig 7A (Basal), 7B (Luminal A)), as well as the MMTRs of those pathways dysregulated across all of the subtypes, were identified. The top 5 MMTRs in Basal-Like unique pathways (SREBF1, ESRRG, ESRRA, RFX2, and SREBF2) differ from those found to be associated with the Luminal A unique pathways (IRF8, OVOL1, THAP1, GATA1, and TFAP2C). Additionally, those MMTRs associated with pathways dysregulated across all subtypes were different from those unique to the Basal-like and Luminal A (HNF4A, ESRRA, HNF4G, RARA, and EP300), with the exception of one (ESRRA), which has been linked to all types of BRCA^57^. Further, the expression levels of these MMTRs cluster out patients based on the subtype of BRCA with which they are associated (Fig 7C), where Basal patients are indicated in black and Luminal A patients are indicated in yellow. The distinct metabolic profiles and ability of MMTRs to accurately distinguish normal Breast Cells from Luminal A and Basal were further confirmed using RNA-sequencing data from 27 different cell lines^58^. First, with the exception of one luminal cell line (JM225CWM), the expression of metabolic genes excellently segregated the luminal, basal and normal breast cell lines (Fig 7D). Secondly, when we clustered the cell lines based on expression levels of the MMTRs of the metabolic pathways uniquely dysregulated in patient samples from Luminal-A subtypes and Basal-like subtypes, they also segregated the cell lines by subtype. These MMTRs offer new explanations for the differences in metabolic reprogramming between different subtypes of BRCA. In total, these findings suggest that the metabolic pipeline has utility in revealing novel understanding with regards to metabolic reprogramming not only across cancers of different tissue origin, but also within heterogeneous cancer populations.

**Figure 7.**
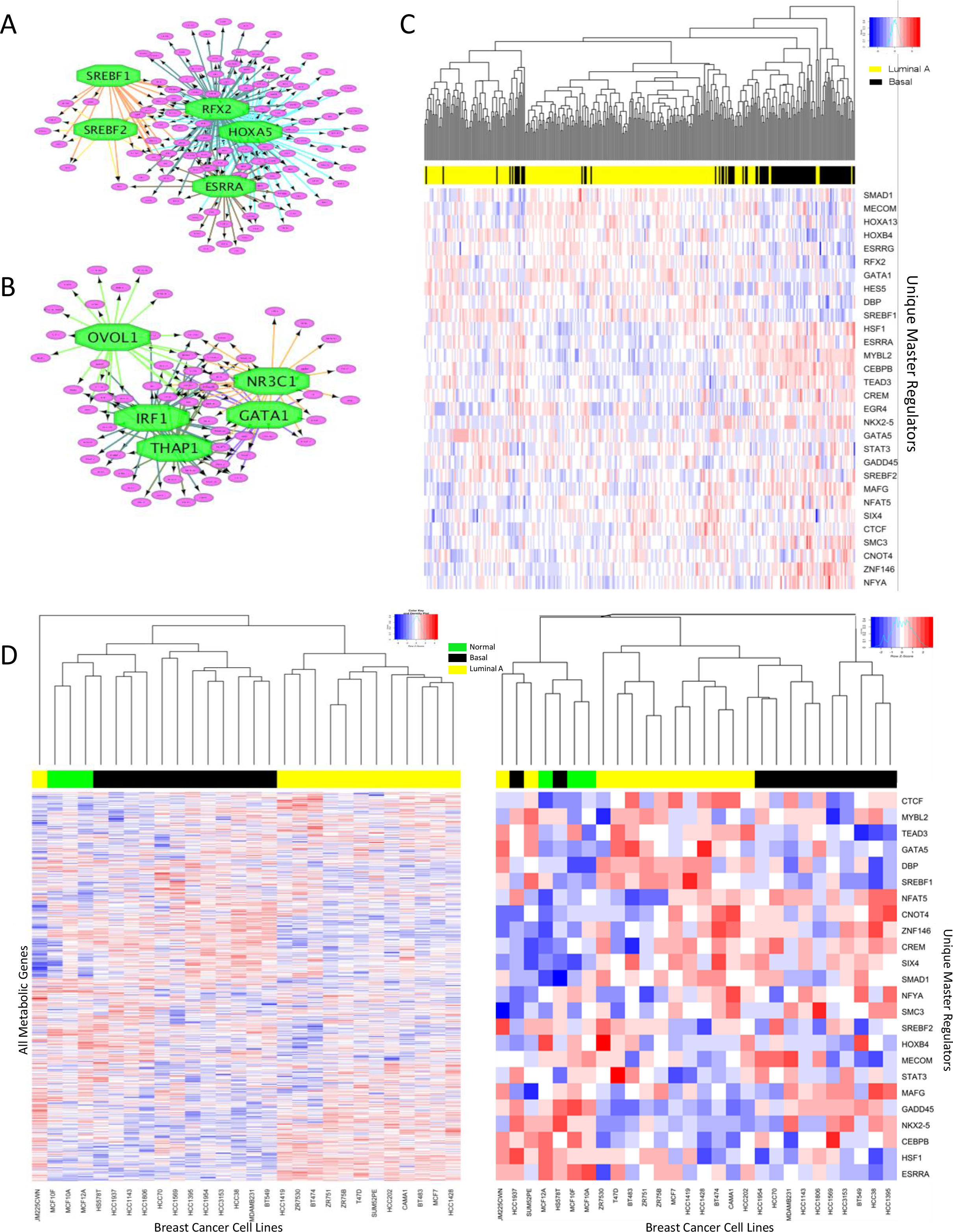
Master Metabolic Transcriptional Regulators (MMTRs) distinguish BRCA molecular subtypes, and BRCA cell lines. (A) MMTR analysis of the pathways unique to Basal-like patients revealed a network of 14 MMTRs, the 5 most highly enriched are shown here. (B) MMTR analysis of the pathways unique to Luminal A patients revealed a network of 15 MMTRs, the 5 most highly enriched are shown here. (C) Unsupervised clustering of Luminal A(yellow) and Basal-Like(black) patients on the expression levels of all MMTRs of their unique pathways create separate clusters of patients. (D) Unsupervised clustering of 27 normal breast (green), basal breast cancer (black) and luminal breast cancer (yellow) patients on all metabolic genes reveals a tight cluster of each of the distinct cell line types. (E) Unsupervised clustering of normal (green), luminal (yellow) and basal-like(black) cell lines on the expression levels of all MMTRs of their unique pathways create separate clusters.

## Discussion

The present study applies an analytical pipeline utilizing transcriptomic information to characterize changes in metabolic pathways associated with cancer. The approach successfully profiled significant metabolic reprogramming in 26 cancer types, revealing both common and unique patterns of disruption. Additionally, this pipeline successfully profiles the metabolic vulnerabilities that distinguish molecular subtypes within the same disease site. These metabolic vulnerabilities represent points of therapeutic leverage in each of these disease sites or subtypes of disease. Furthermore, cancer specific expression patterns are linked to the identification of metabolic master transcriptional regulators (MMTRs), which provide putative mechanistic insight into observed metabolic profiles. MMTRs associated with target expression, genetic drivers and clinically relevant molecular signatures in cancer cohorts, suggesting them as key regulators of metabolic reprogramming in cancer. Additionally, MMTRs can be used as a predictive biomarker of metabolism-targeting drug response, as well as a means of segregating molecular subtypes within a disease site (Fig 7).

While metabolite levels are the final output, and therefore the most sensitive gauge of metabolic activity, there is a lack of pan-metabolic data across disease sites. Due to the dynamic nature of connected metabolomics and transcriptomic changes, generating models of metabolism based on transcriptomics is complex. To better understand metabolic flux across types of cancer and even among sub-types within a particular cancer type, more unbiased metabolomics studies need to be conducted to more fully appreciate the role of metabolic reprogramming in cancer initiation, progression, and prognosis^59^.

A transcriptomic metabolic view of TCGA^12^ data, where metabolic pathways scores are generated, yields insights that cannot be acquired through differentially expressed genes analysis. This pipeline expands on DEG analysis alone by not only looking at the magnitude of the changes that occur in a particular gene, but also, how meaningful that change is to the disease site. Combining the log fold-change and the adjusted p-value into a single score allows us to scale the importance of each gene expression change within metabolic pathways. The bootstrapping approach accounts for background changes in a way that identifies patterns that would be expected simply by chance, allowing for the elucidation of relevant transcriptional metabolic changes. This is an important aspect of our pipeline, as it helps to account for the varying degree of tumor associated transcriptional drift across cancer types, as well as for tissue procurement error and/or contaminating cell types that may be associated with cohort samples at tissue specific rates. Furthermore, mapping these genes, with their direction and magnitude of change (up or downregulation) onto the pathway allows for determination of patterns that indicate a convergence of effect on key metabolites.

A high degree of correlation between population pools from different transcriptomics platforms (RNA-sequencing vs. microarrays) further demonstrates the robustness of this approach (Supp Fig 1). These datasets implicate many of the same pathways as being highly dysregulated in PRAD^39^, LUAD^40^ and BRCA^41^ as compared to the normal matched tissues, and to largely the same relative magnitudes (Supp Fig 1). This confirmation in 3 separate populations of patients on a different transcriptomic platform reveals we are highlighting biologically relevant metabolic pathway dysregulation, and our scoring approach is highly robust.

An important test of the validity of the metabolic pathway scores generated from this approach is to model metabolic circuits, predict expected alterations in metabolite pools, and compare that with published studies that examine those metabolites. For example, in Figure 3, we explore the Pentose Glucuronate Intercoversion pathway, the dysregulation of which is well-known and almost exclusively studied in LIHC^42,43^. However, we suggest there are cancers that dysregulate this pathway to an even greater degree, including LUAD, in which we find significant upregulation of many genes within the pathway. Unbiased metabolomics studies comparing LUAD with normal lung tissue identified significant elevation of UDP-D-Glucuronate^44^, which is the predicted result of the changes in expression level of enzymes in this pathway. As shown in Figure 3C the significant gene expression changes in LUAD patient samples would be expected to divert metabolites towards the production of UDP-D-Glucuronate. This metabolite is responsible for the downstream production of UDP-Glucose and a-D-glucose-1P, which feed into several other pathways, fueling polysaccharide biosynthesis and glucosiduronide production^60^. The connection to these pathways are relevant to the disease because they support a high rate of nucleic acid synthesis, and provide NADPH for both the synthesis of fatty acids and cell survival under stress conditions.^61,62^ Interestingly, LUSC upregulates this pathway more so than LUAD in regards to magnitude of transcriptional change. However, little metabolomic data exists in the context of this disease site. Nevertheless, our metabolic pathways scores implicate this pathway as being at least equally important in lung squamous cell carcinoma and adenocarcinoma.

Perhaps not surprisingly, we found some metabolic pathways that are dysregulated in most or all types of cancer, such as the glycolytic and gluconeogenesis pathways, as well as pathways that have a highly restricted pattern of dysregulation like polyamine metabolism in prostate cancer (Fig 4A). The global upregulation of genes within the pathway (Fig 4C) involved both the generation of acetylated polyamines and flux through biosynthesis utilizing ornithine and sadenosylmethionine to eventually produce spermine and spermidine. It is well-established that this pathway is highly active in normal prostate, due to high rates of secretion of acetylated polyamines into prostatic fluid, and further enhanced in prostate cancer.^47,48^. The nearly complete upregulation of the biosynthetic and catabolic enzymes in PRAD is striking, and the idea of increased metabolic flux is supported by metabolomics^49^. Additionally, the therapeutic targeting of this pathway with BENSpm is highly effective and selective, highlighting the utility of the metabolic pipeline to determine metabolic pathways of interest for pharmacologic intervention. Differences in metabolic pathway dysregulation and therapeutic targeting may be attributed to the identification of MMTRs. We identified a set of MMTRs for the genes in the polyamine metabolic pathway whose expression highly and positively correlated with the significant upregulation of those genes. In contrast, the polyamine pathway is down regulated in KICH and exhibited weaker and non-significant correlations between expression of MMTRs and the genes within the pathway. Further, association of MMTRs with common cancer type specific mutations may indicate differences in metabolic reprogramming in specific patient populations based upon co-occurrence or mutual exclusivity. For example, TMPRSS2-ERG fusion in PRAD, which is one of the most frequently occurring mutations, is mutually exclusive with GTF2B overexpression, a highly enriched MMTR. Interestingly, there is a significant amount of overlap amongst ERG and GTF2B binding sites, and in PRAD ChIP-sequencing data, ERG peaks have been identified in three of the polyamine genes, potentially explaining their mutual exclusion. (Supp Fig 4) This therefore highlights the potential for different genetic drivers of disease to cooperate with altered expression of MMTRs in order to drive specific patterns of metabolic reprogramming.

Also of interest was the identification of subsets of patients within breast cancer that exhibit very different patterns of metabolic reprograming. Understanding the metabolic profiles of different molecular subtypes is important in disease sites like BRCA, where the different subtypes have distinct treatment regimens and outcomes (Fig 6). Pooled analysis, treating all subtypes as one cohort leads to an understanding of BRCA as a whole (Fig 6A, 6B), but fails to capture the subtypes that are driving the overall analysis. For example, the Basal-Like subtype, as defined by the PAM50^53,54^ and also known as Triple Negative Breast Cancer (TNBC), clusters very well based on metabolic genes and exhibits more highly dysregulated metabolic pathways (Fig 6C). We effectively targeted this metabolic difference with drugs that target the Citric Acid Cycle (i.e. Metformin) (Fig 6G), and drugs that target the Homocysteine Biosynthetic Pathway (i.e. Sulfasalazine)(Fig 6H). We found that, in both cases, the basal-like cell lines are more sensitive than the Luminal A like cell lines, in agreement with the fact that these pathways were uniquely dysregulated in the basal-like patient data. This again highlights the ability of the metabolic pipeline to emphasize metabolic pathways of interest that distinguish molecular subtypes within a single cancer, and provides selective therapeutic targets. The identification of MMTRs driving the differences in metabolic dysregulation between the luminal A and basal subtypes results in distinct clustering of the Basal and Luminal A subtypes when looking at the expression of those MMTRs (Fig 7). This, therefore, offers novel opportunities to understand how and why metabolic pathways differ between tissues sites and molecular subtypes, and offers new potential targets.

Overall, we provide a novel approach to utilizing transcriptomic data to identify metabolic dysregulation. While it is clear that metabolic regulation occurs to a great extent at the post-transcriptional and even post-translational levels, this transcriptome based approach provides novel insights. These insights and transcriptomic analyses will need to be combined with proteomic and metabolomics data to develop a comprehensive picture of metabolic dysregulation in cancer. Nevertheless, the high level of correlation observed between studies in separate cohorts of patients with the same disease, and the metabolomics support we have in some pathways give us confidence that this method is highly informative pan-cancer. This analytical pipeline can be applied to any transcriptome wide data to infer patterns of metabolic reprogramming, in any disease settings.

## Methods

### Pan-Cancer Differentially Expressed Gene Analysis

The results published here are in whole based upon data generated by The Cancer Genome Atlas (TCGA)^12^ Research Network: http://cancergenome.nih.gov/. Firehose^63^, a web portal site that has been developed by the Broad Institute, aiming to deliver automated analyses of the TCGA data to general users, was utilized to download the preprocessed, Level 3, RSEM transcriptomic data. Gene expression data were analyzed using Bioconductor 3.1 (http://bioconductor.org)^64^, running on R 3.1.3. RNA-sequencing RSEM counts were processed to remove genes lacking expression in more than 80% of samples. To identify differentially expressed genes, primary tumor samples (samples ending in “.01”) were compared to their matched normal tissues (samples ending in “.11”), in their respective tissues. Scale normalization and moderated student t tests were performed using empirical Bayes statistics in the “Limma”^65^ package. The resulting P values were adjusted for multiple testing using the false discovery rate (FDR) Benjamini and Hochberg corrections method.

### Code Availability

Code is available upon request to the corresponding author.

### Pathway Score

Gene and pathway scores were calculated in R 3.1.3. Differentially expressed gene lists for each cancer site were used to assign individual gene scores. Gene scores were designated by taking the absolute value of the logFC multiplied by the – log(adjusted P value):

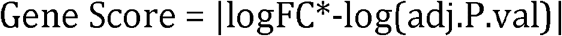

Metabolic pathways were then downloaded from the Kyoto Encyclopedia of Genes and Genomes (KEGG)^38^ were downloaded. Genes from each of the 114 pathways are reported in the Supplement (**Supplementary Table 2**). Pathway scores were then calculated by summing the gene scores for all genes within each of the pathways and dividing by the square root of the sample size for that particular tissue, to account for sample size effects in different cancer sites:

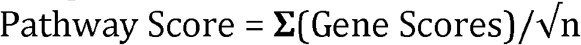

All pathway scores were then exported into a table, to determine statistical significance of each score (**Supplementary Table 3**). Pathways were clustered into 10 major categories based upon KEGG classifications.

### Bootstrapping for Pathway Score Statistical Significance

Bootstrapping^66^ is a technique based on random sampling with replacement. Using R 3.1.3, pathway scores were randomly generated 100,000 times per pathway, based on the number of genes in the pathway, and plotted into a distribution. The scores for each of those pathways were then plotted against the distribution and a P value was calculated based on where that score lies within the distribution of scores (**Supplementary Figure 5**). Using all p-values, the pathway score table (**Supplementary Table 4**), was adjusted to only include those scores that were considered to be statistically significant. All other values were replaced with “0” (**Supplementary Table 5**).

### Pathway Scores Heatmap

Bootstrapped pathway scores were utilized to create pathway score heatmaps in R 3.1.3, constructed using the “Gplots” and “Heatmap.2” packages in R. Data was scaled and Euclidian distances and hierarchical clustering were applied using the “h.clust” function. All 0 values (non-significant pathway scores) are represented as gray. For specific pathway heatmaps, at the gene level, fold change values from the initial Limma output for each cancer type, was utilized. Data was scaled using a min to max calculation:

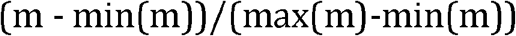

Once again Euclidian distances and hierarchical clustering was applied. Heatmaps of the significantly differentially expressed genes are represented by blue (negative) or red (positive), and non-significantly differentially expressed genes are represented as gray.

### Pathway Maps

Pathway maps were generated using Cytoscape^67^ software, and specifically the VizMapper functions. Pathway maps were based on existing pathway maps in KEGG^38^. Limma output for differentially expressed gene analysis was utilized to direct shading of genes within the pathway: red (positive fold change, statistically significant), blue (negative fold change, statistically significant), or gray (non-statistically significant), for individual cancer sites.

### Dose Response Cell Viability

PC3 and DU145 cells were obtained from ATCC [Manassas CA]. MDA-MB-231 cells were provided by Dr. John Ebos, Ph.D. [Department of Cancer Genetics, Genomics and Development, Roswell Park Comprehensive Cancer Center (RPCCC), Buffalo, NY]. 786-O and ACHN cells were provided by Dr. Eric Kauffman, M.D. [Department of Medicine, Roswell Park Comprehensive Cancer Center (RPCCC), Buffalo, NY]. HS578T cells were provided by Dr. Mikhail Nikiforov, Ph.D. [Department of Cell Stress Biology, Roswell Park Comprehensive Cancer Center (RPCCC), Buffalo, NY]. MCF7 and T47D cells were provided by Dr. Katerina Gurova, M.D., Ph.D. [Department of Cell Stress Biology, Roswell Park Comprehensive Cancer Center (RPCCC), Buffalo, NY]. All prostate cancer cells (DU145 and PC3) were maintained in RPMI 1640 medium supplemented with 10% FBS, and 1% antibiotics. Other cells were maintained in DMEM with 10% FBS and 1% antibiotics. Metformin and Sulfasalazine were obtained from Sigma and N1, N^11^-bis(ethyl)norspermine (BENSpm) was purchased from Synthesis Med Chem [Shanghai, China]. Cells were seeded in 96-well plates at 3000 cells/well on day 0. They then underwent either 48 hours (BENSpm and Sulfasalazine) or 72 hours (Metformin) of treatment. Resazurin (Sigma) was then added to each well and allowed to incubate for 2 hours at 37°C. The plates were then read on a spectrophotometer by excitation at 570 nM and reading of the fluorescence at 600 nM. Dose-response curves were then plotted using Prism GraphPad 7.

### Master Metabolic Transcriptional Regulator Analysis

In order to characterize regulatory networks, we used iRegulon^51^, a Java add-on in Cytoscape, to identify MMTRs. In this approach, we use a large collection of transcription factor (TF) motifs (9713 motifs for 1191 TFs) and a large collection of ChIP-seq tracks (1120 tracks for 246 TFs). This method relies on a ranking-and-recovery system where all genes of the human genome (hg19) are scored by a motif discovery step integrating the clustering of binding sites within *cis*-regulatory modules (CRMs), the potential conservation of CRMs across 10 vertebrate genomes, and the potential distal location of CRMs upstream or downstream of the transcription start site (TSS+/−10 kb). The recovery step calculates the TF enrichment for each set of genes, input for each of the individual analyses, leading to the prediction of the TFs and their putative direct target genes which exist in the input lists. This method optimizes the association of TFs to motifs using both direct annotations and predictions of TF orthologs and motif similarity.

### MMTR Correlation Analysis

Correlation values between MMTRs and all expressed genes were derived in R 3.1.3, constructed using the “cor” function across all patients in both TCGA-PRAD and KICH cohorts. The empirical cumulative distribution function for each complete MMTR correlation profile (background) was determined via the “ecdf” function, and similarly for the MMTR correlation profiles against Polyamine Biosynthetic genes only. Significant shift in distributions between MMTR/background and MMTR/Polyamine Biosynthetic gene correlations was assessed by Kolmogorov-Smirnov test.

### Common Mutation Analysis

Using cBioPortal (http://www.cbioportal.org), patients in the PRAD cohort were queried for either co-occurent relationships or mutually exclusive relationships between the list of most commonly occurring mutations in PRAD and the 4 MMTRs in question. CBioPortal is a publically available database, which based on all of the genomic data available for PRAD constructs a list of the most commonly occurring mutations. Additionally, it calculates the significance of co-occurrence or mutual exclusivity, based upon the Mutual Exclusivity Modules (MEMo)^65^. MEMo is a method that searches and identifies modules based upon: (1) genes recurrently altered across a set of tumor samples; (2) genes known to or likely to participate in the same biological process; and (3) alteration events within the modules are mutually exclusive. Using this information, it then integrates multiple data types and maps genomic alterations to biological pathways and uses a statistical model that predicts the number of alterations both per gene and per sample.

### GTF2B and ERG Binding Site Overlap

ChIP-Sequencing BED files were downloaded from the Cistrome database^66^, corresponding with 3 different studies. To confirm overlap between GTF2B and ERG: K562 Erythroblast; Bone Marrow Untreated from the Martens JH, et al. (2012)^67^ study and GTF2B K562 Erythroblast; Bone Marrow Untreated from the Pope BD, et al. (2014)^68^ study were downloaded. GenomicRange was then used to determine the overlap between these peaks in the same line. Then, to determine ERG peaks in Polyamine Biosynthetic genes in prostate cancer cell lines specifically, the VCaP; Epithelium; Prostate ERG non-treated data from Sharma NL, et al. (2014)^69^ was downloaded and imported into the Interactive Genome Viewer (IGV)^70^ to visualize peaks in these genes. GTF2B ChIP-sequencing data was not available for GTF2B in prostate cancer.

### PAM50 BRCA Analysis

The PAM50 is a method that has been previously described in the literature^53,54^. The PAM50 classification of tumors within the TCGA cohort was obtained^54^. This classification was then used to stratify patients into 4 major groups: Basal-Like, HER2-expressing, Luminal A, and Luminal B. Any patients with no classification within this file were removed from the analyses and all data was preprocessed as outlined in the *Pan-Cancer Differentially Expressed Gene Analysis* section. Post-normalization, all tumors still included underwent unsupervised clustering based on the expression of all metabolic genes within the cohort. Additionally, comparisons of patients within each cohort were then made with normal tissues to obtain Pathway Scores, as previously described, for each of the patient cohorts. MMTR analysis was then performed to determine MMTRs of uniquely dysregulated pathways within the Basal-Like and Luminal A subtypes. Patients from each of those subtypes then underwent unsupervised hierarchical clustering based on the gene expression of those MMTRs of uniquely dysregulated pathways.

### BRCA Cell Line Analysis

RNA-sequencing data was obtained for 27 different normal breast, luminal breast cancer and basal breast cancer cell lines^58^. All data was preprocessed as outlined in the *Pan-Cancer Differentially Expressed Gene Analysis* section. Post-normalization, all tumors still included underwent unsupervised clustering based on the expression of all metabolic genes within the cohort. Additionally, comparisons of the cell lines from each of those subtypes then underwent unsupervised hierarchical clustering based on the gene expression of those MMTRs of uniquely dysregulated pathways, from the patient data.

## Author Contributions

S.R.R. contributed to the experimental design, experiments, analysis, and writing. M.D.L. contributed to the experimental design, analysis and writing. H.C.A. contributed to the experimental design and writing. A.M.R. contributed to the experimental design and writing. K.H.E contributed to the experimental design. D.J.S. contributed to the experimental design and writing.

